# The shifting importance of abiotic and biotic factors across the life cycles of wild pollinators

**DOI:** 10.1101/2022.04.25.489447

**Authors:** Jane E. Ogilvie, Paul J. CaraDonna

## Abstract

1. Organisms living in seasonal environments are exposed to different environmental conditions as they transition from one life stage to the next across their life cycle. How different life stages respond to these varying conditions is a fundamental aspect of biology and is critical for understanding how organisms will respond to environmental change.
2. Despite the importance of animal pollinators, we lack a basic understanding of the influence of different environmental factors across their life cycles. We investigated the relative importance of climate conditions, food availability, and previous life stage abundance in a community of wild bumble bee species, asking: how do these three factors influence bee abundance at each life stage?
3. We used a 7-year dataset to examine the importance of environmental conditions on the abundance of life stages in seven wild bumble bee species. We monitored climate conditions, the abundance of floral resources, and abundances of bees in each life stage across the active colony life cycle in a highly seasonal subalpine ecosystem in the Colorado Rocky Mountains, USA.
4. Bee abundance at different life stages responded to environmental conditions in a consistent manner across the seven species. The survival and recruitment stage of the life cycle (assessed as abundance of overwintered queens) responded negatively to longer winters; the growth stage (assessed as abundance of workers) responded positively to floral resource availability; and the reproductive stage (assessed as abundance of males) was positively related to the abundance of the previous life stage (workers).
5. Our long-term examination of annual bumble bees reveals a consistent set of responses in the abundance of each life stage to climate conditions, floral resource availability, and previous life stage abundance. Across species, these three factors each influenced a distinct life stage, illustrating how their relative importance can shift throughout the life cycle. The life-cycle approach we have taken with wild bumble bees highlights that important details about pollinator demography are overlooked without considering life stage-specific responses. Ultimately, it is these life-stage specific responses that shape population outcomes, not only for animal pollinators, but for many organisms living in seasonal environments.

## 1 INTRODUCTION

Climate conditions and food resources are two key dimensions of an organism’s environment that influence survival, growth, and reproduction. How an organism responds to these abiotic and biotic factors throughout its life cycle will shape population dynamics (Caswell, 2001; White, 2008; Ehrlén et al., 2016). In seasonal environments, variation in the abiotic and biotic environment may be particularly important because different life transitions (e.g., recruitment, growth, reproduction) can occur under different conditions (Elton, 1927; Fretwell, 1972; Radchuk et al., 2013). If the abundances of successive life stages covary, then factors affecting the abundance of one life stage can influence the abundance of the next, with implications for population dynamics (Caswell, 2001). Understanding the relative importance of abiotic and biotic factors across an organism’s life cycle is particularly important in the context of anthropogenic climate change (e.g., Radchuk et al., 2013; Oro, 2013), because climate change not only alters the abiotic environment, but it can also alter the biotic environment by influencing resource availability and species interactions.

We currently know relatively little about the factors driving the demography and thus population dynamics of wild pollinators (Roulston & Goodell, 2011; but see Boggs & Inouye, 2012; Crone & Williams, 2016; Wong & Forrest, 2021)—despite the critical ecosystem services pollinators provide (Roubik, 1995; Kremen et al., 2002; Klein et al., 2007; Garibaldi et al., 2013) and their widely observed declines (Goulson et al., 2015; Potts et al., 2016). The abundance of different pollinator life stages can be influenced by climate conditions experienced, the availability of floral food resources, and the success (as assessed e.g., by abundance) of previous life stages. Indeed, two major factors underlying observed global pollinator declines are reduced food resources (Potts et al., 2016; Woodard & Jha, 2017) and changing climate conditions (Ogilvie et al., 2017; CaraDonna et al., 2018; Soroye et al., 2020; Crossley et al., 2021). Although these factors define basic requirements for pollinators, we lack a basic understanding of how their influence might vary across different life stages, especially in wild ecosystems.

Here we investigate the relative importance of climate conditions, food availability, and the abundance of previous life stages across the life cycles of seven wild bumble bee species that coexist within the same ecological community. Specifically, we ask: how do these three factors influence the abundance of each bumble bee life stage? To address this question, we used a unique long-term study where we have monitored floral resource abundance, climate conditions, and bee abundance across the active bumble bee colony life cycle over seven years in a subalpine ecosystem in the Colorado Rocky Mountains.

Bumble bees (*Bombus* spp.) are eusocial animals with an annual colony life cycle that has distinct life stages corresponding to general demographic parameters: survival and recruitment (reflected in the abundance overwintered queens); growth and size (reflected in the abundance of workers); and reproduction (reflected in the abundance of males and autumn queens produced near the end of the active colony season) (Fig. 1). Although the abundance of each life stage can be influenced by climate, floral resources, and the abundance of the previous life stage, it is unlikely that each life stage will respond to these factors similarly because each performs a different function, is active during a different time of the year, and therefore experiences a different abiotic and biotic environment. For example, the abundance of overwintered queens observed in spring is surely sensitive to climate conditions experienced during winter diapause, whereas the abundance of workers and reproductive output may be more sensitive to food resources and climate conditions experienced during summer. By examining wild bumble bee populations across their life cycles over multiple years, our work helps to fill an important knowledge gap in our basic understanding of pollinator populations— especially in the context of climate change and pollinator declines.

**Figure 1.**
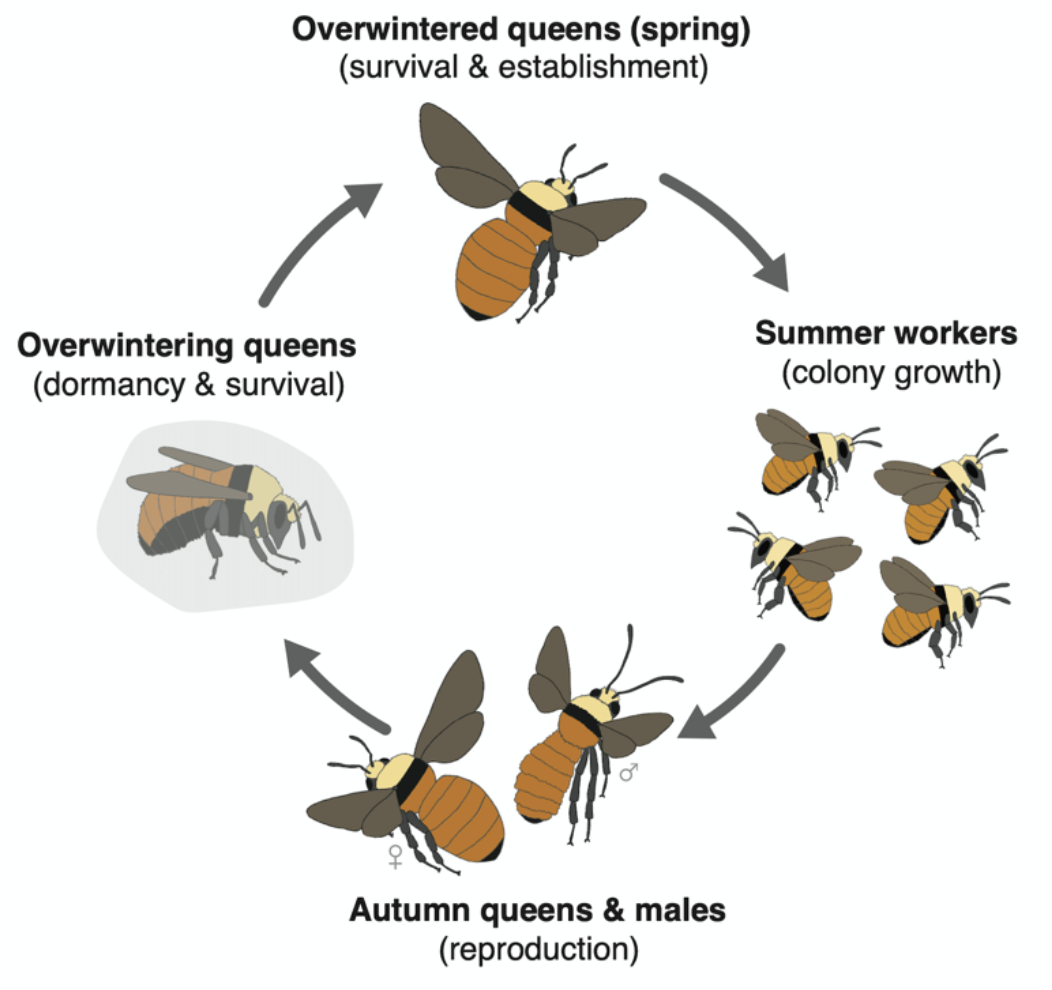
Annual life cycle of a bumble bee colony illustrating each life stage and its corresponding demographic parameter.

## 2 MATERIALS AND METHODS

### 2.1 Study system

We monitored floral resources, climate conditions, and the abundance of seven wild bumble bee species across seven summers (2015-2021) in subalpine habitats near the Rocky Mountain Biological Laboratory (RMBL) in Gothic, Colorado, USA (2900 m a.s.l.). The short season of most biological activity in this system begins when snow melts in spring (April–June) and ends when temperatures cool in autumn (September–October). Thus, the study area typifies high-elevation and high-latitude locations with extreme seasonal climates, which are experiencing especially rapid anthropogenic climate change (IPCC, 2007; Nogués-Bravo et al., 2007).

We chose six study sites spread over 8 km within the East River Valley near the RMBL, each separated from the others by at least 1 km (from edge to edge), and each approximately 500 m in diameter (196,349 m^2^)—a scale that reflects typical bumble bee foraging distances observed in Colorado montane meadows (Elliott, 2009; Geib et al., 2015). Mark-recapture data from two years of study at these sites provides evidence that our focal bumble bee species do not travel between them (no site transfers from 2,012 marked bees). Each site includes the three major habitats where bumble bees are observed foraging or searching for nests: wet meadow, dry meadow, and aspen forest. Across the six sites there are 153 non-graminoid herbaceous and shrub flowering plant species from 40 plant families, 96 species of which we have observed bumble bees to visit, although 99% of bumble bee visits are to 48 plant species. Five of the plant species visited by bumble bees are introduced (the common *Taraxacum officinale* and *Trifolium repens*, and rarer *Trifolium pratense, Linaria vulgaris*, and *Cirsium arvense*). Non-native bees, including the honey bee, *Apis mellifera*, are absent in this area.

### 2.2 Bumble bees

Our observations focused on the seven most common bumble bee species: *Bombus appositus, B. bifarius, B. flavifrons, B. insularis, B. mixtus, B. occidentalis*, and *B. rufocinctus*. *Bombus insularis* is a nest parasite of other bumble bees and thus does not have workers. These seven species coexist in our system and belong to the same ecological community (Pyke, 1982).

Globally, bumble bees are a rich group of approximately 260 species in mostly temperate and montane ecosystems, though many species are experiencing contractions in their distribution and relative abundance (Cameron & Sadd, 2020). Bumble bee species, including those within our study assemblage, vary in life history traits, including diet breadth, phenology, and body size—all of which may influence their response to changes in the abiotic and biotic environment (Ogilvie & Forrest, 2017; Martinet et al., 2021). Bumble bees are quintessential generalist foragers (Kleijn & Raemakers, 2008; Wood et al., 2021; Ogilvie & CaraDonna, in review). Their floral resource needs extend across most of the growing season as they transition from one life stage to the next, and each life stage is exposed to a new set of abiotic conditions (Fig. 1). In spring, overwintered queens that mated the previous autumn and have survived winter diapause emerge to search for nests and establish colonies. The abundance of overwintered queens in spring should be related to floral resources and colony reproductive output during the previous growing season when they were produced (Inari et al., 2012; Carvell et al., 2017), as well as the winter conditions experienced during diapause (Vesterlund & Sorvari, 2014). If overwintered queens are successful in colony establishment, their colonies grow in the number of foraging workers over the summer and eventually produce reproductive bees toward the end of the season (autumn queens and males). Both colony growth and reproductive output may be influenced by the availability of floral resources during summer (Crone & Williams, 2016), prevailing climate conditions (Iserbyt & Rasmont, 2012), and the abundance of the previous life stage (Inari et al., 2012).

### 2.3 Quantifying bumble bee abundance

We monitored the abundance of bumble bees of each species and life stage at weekly intervals across the entire season of above-ground activity (16–22 weeks, late April to late September). During each weekly census we systematically searched the three habitats at each study site (dry meadow, wet meadow, aspen forest) for 20 minutes each, recording all bumble bees seen. For each bee we recorded: (i) species identity; (ii) life stage (overwintered queen, worker, male); (iii) whether the bee was foraging, searching for a nest, or both; and (iv) the plant species visited if the bee was foraging. Bees were typically identified to species on the fly based on distinctive colour patterns and body size following Williams et al. (2014), but if the identity was uncertain, we netted the bee for closer inspection and quickly released it. These methods yielded estimates for the abundance of each life stage for each of the seven species in each of the six sites and seven years. For each species and life stage we summed the abundances of bees seen in each weekly observation (number of bees per hour for all weeks observed). We standardised bee counts in each week to express them as number of bees per hour, to accommodate rare instances where our weekly censuses at a given site were longer or shorter than one hour.

### 2.4 Quantifying the effect of previous life stage

To understand the extent to which demographic responses are linked across the bumble bee life cycle, we examined how each life stage is influenced by the abundance of the previous life stage. For overwintered queens this was males observed in the previous season (t-1); for workers this was overwintered queens in the current season; and for males this was workers in the current season (Fig. 1). For overwintered queens, we also considered the abundance of workers in the previous season (t-1), since this is the most common life stage measured in other bumble bee population studies and it has been shown to relate the production of both males and new queens (e.g., Pelletier & McNeil, 2003; Crone & Williams, 2016; Spiesman et al., 2017); however, we note that our results are qualitatively similar when we use either workers or males as a predictor of overwintered queen abundance. We did not include autumn queens as a distinct life stage or include them in reproductive output because the observation of new, autumn queens is rare in our study system. Thus, reproductive output is represented by male abundance.

### 2.5 Quantifying floral resource abundance

In the same weekly censuses, we also recorded floral abundance. At each site we counted the flowers of plant species visited by bumble bees in 15 permanent 20 x 0.5 m belt transects, five distributed throughout each of the three habitats (i.e., 50 m^2^ of transect area per habitat, 150 m^2^ total transect area per site). We counted individual flowers of most species, but instead counted the number of flowering stalks for *Castilleja* spp., *Eriogonum* spp., *Valeriana occidental*, and *Haeckelia floribunda* as well as members of the Apiaceae and Lamiaceae; similarly, we counted the number of capitula for all Asteraceae and the number of catkins for *Salix* spp.

We quantified abundance of floral resources for bumble bees by first generating a list of all flower species visited by each species and life stage across the seven years. This list was used to create a floral preference index by dividing the number of visits to each species by the number to the most-visited species (following Pleasants, 1981). Rescaling the flower counts by this index prevents rarely-visited species from representing an inappropriate share of a bee diet. For example, *Bombus bifarius* rarely visits the early spring *Claytonia lanceolata*, so this species should be scored as a minor element of their diet even though the flowers are abundant.

The rescaled flower counts then allowed us to estimate those resources that had the potential to influence the abundance of each bumble bee life stage. For overwintered queens (produced toward the end of each growing season but observed the following spring; Fig. 1), our estimate of resource abundance includes all flowers visited across the entire previous growing season (t-1). For workers, our estimate includes all flowers visited from the start of the growing season through the end of observed worker activity. For males, our estimate includes all flowers visited from the start of the growing season through the end of observed male activity.

### 2.6 Quantifying climate conditions

We selected climate variables that are hypothesised to influence the abundance of each life stage and that each life stage directly experiences (Iserbyt & Rasmont, 2012; Ogilvie et al., 2017). For overwintered queens, our climate variable is spring snowmelt date—the date when bare ground appears and emergence from winter hibernacula is possible. In our system, earlier spring snowmelt indicates a shorter winter with less snowpack. For workers and males, our climate variable is the ratio of temperature to precipitation (mean daily maximum temperature divided by the sum of precipitation) which captures variation in both summer temperature and precipitation when bees are active. For this ratio, lower values indicate cooler and wetter conditions, whereas higher values indicate warmer and drier conditions. For workers, the temperature:precipitation ratio includes June and July conditions, and for males, it includes July and August conditions—these are times when the two life stages are mostly active. Climate conditions were measured by local resident billy barr at a central site at the RMBL, 0.5–4.5 km from the six study sites. Snowmelt date was the day of year that a permanent 5 x 5 m plot was first free of snow, while temperature minima and maxima (in °C) and precipitation (in mm) were measured daily using a Davis Instruments temperature sensor and a standard US National Weather Service rain gauge, respectively.

### 2.7 Data analysis

We analysed how climate conditions, floral resources, and previous life stage were related to bee abundance by constructing generalised linear mixed models with each of these factors as additive predictor variables for each bumble bee species and life stage. Study site was included in all models as a random intercept term. The bee counts are overdispersed, so we used a negative binomial error distribution. Analyses were conducted in a Bayesian statistical framework (using the ‘stan_glmer.nb’ function). All models used weakly informative priors and were created in the Stan computational framework (http://mc-stan.org/) accessed with the *rstanarm* package in R (Goodrich et al., 2020). Model estimates (i.e., the effect of each factor) from our negative binomial models are expressed as the change in the log count of bees per 1 unit change in the predictor variable (except for flowers, which for simplicity are shown as the change in the log count of bees per 100 flowers). Bayesian R^2^ values were calculated for each full model using residual variance following Gelman et al., (2019). The Bayesian statistical framework places an emphasis on the uncertainty of predictions produced from a given model (i.e., the parameter estimates) as opposed to a single point estimate (Ellison, 2004). Evidence for a hypothesis can be expressed as the probability of a given parameter estimate (i.e., the posterior distribution of the model estimates; Ellison, 2004). We consider strong evidence for a non-zero model parameter estimate when the 89% Bayesian credible interval from the posterior distribution does not include zero; in other words, this means that at a minimum 89% of the predictions from a model support the hypothesis of a non-zero effect. All analyses were conducted in R version 4.0.2 (R Core Team, 2020).

## 3 RESULTS

Seven years of season-long censuses yielded counts of 28,639 bees, 1,004,241 flowers, and 700 unique interactions between flowers and life stages of the different bumble bee species. Over this time, we detected no clear directional trends in the abundance of bumble bee species and life stages (Fig. 2). Climate conditions varied considerably from 2015 to 2021: spring snowmelt dates varied by one month (May 5–June 6); the summer average high temperature spanned almost 3.0°C (19.9–22.7°C, June through August); summer precipitation varied more than three-fold (78–272 mm, June through August); and the ratio of temperature:precipitation varied five-fold (0.1–0.5). Likewise, floral resource abundance for all flowers visited by bumble bees varied seven-fold across years and study sites (959–6966 flowers/m^2^, summed over each season).

**Figure 2.**
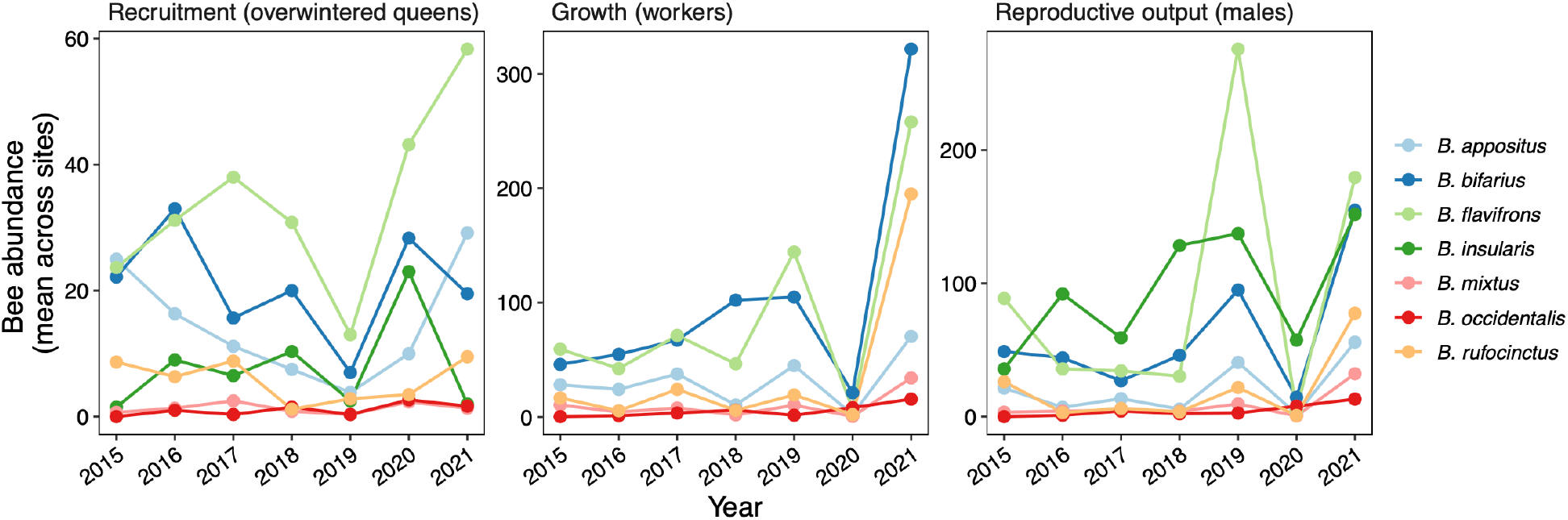
Mean abundance of seven bumble bee species (*Bombus* spp.) and life stages over 7 years. Bee abundances are averaged across the six study sites in each year. Note the different abundance scales for each life stage; *B. insularis* is a bumble bee nest parasite without a worker life stage.

Across bumble bee species, our full additive models relating the abundance of different life stages to putative environmental drivers explained 39–60% of the variation for overwintered queens; 42–60% for workers; and 40–70% for males (range of values represents median Bayesian R^2^ values across species; Fig. 3; Table S1). Life stages responded to variation in climate, floral resources, and previous life stage in some consistent ways across species, but most species also exhibited some idiosyncratic responses (Fig. 3; Fig. 4; Figs. S1–5). We can illustrate both patterns using the examples of the two most common species, *B. bifarius* and *B. flavifrons* (Fig. 4). As was generally the case across species, the abundance of overwintered queens responded negatively to winter climate conditions; for species with evidence of a non-zero response, on average, we observed a 2.3-fold decline in bee abundance across the observed range of snowmelt dates. The abundance of workers responded positively to the availability of summer floral resources; for those species, we observed a 3.88-fold decline across the range of observed floral resources. Lastly, the abundance of males responded positively to the abundance of the previous worker life stage; for those species, we observed a 4.89-fold decrease in the abundance of bees across the observed range of workers.

**Figure 3.**
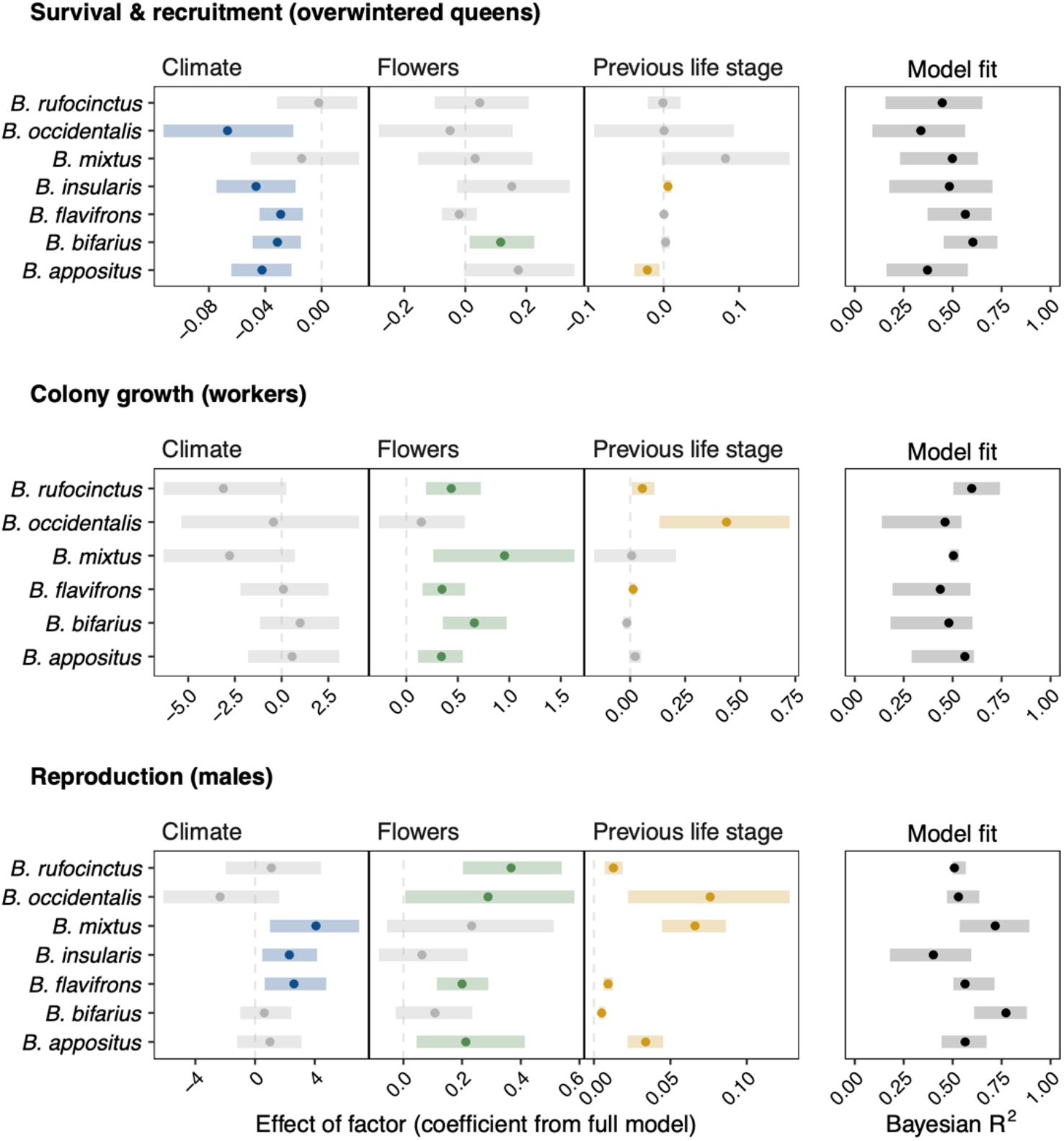
The effect of climate conditions, floral abundance, and previous life stage on the number of bees in different life stages in the seven bumble bee species. Dots represent the median estimate (i.e., effect) of the posterior distribution from negative binomial mixed models including all three factors; error bars represent 89% Bayesian credible intervals. Coloured dots and error bars represent model factors with evidence in support of a non-zero effect (e.g., 89% credible interval does not include zero). The effect of each factor (coefficients from the full model) can be interpreted as follows: for every one-unit change in the predictor variable, the expected log count of bees changes by the corresponding coefficient estimate (with the other predictor variables in the model held constant); for flowers, coefficients are shown as change for every 100 flowers. Model fits represent the median and 89% credible interval for Bayesian R^2^ values. The climate variable for overwintered queens is spring snowmelt date; for workers is the June and July temperature:precipitation ratio; and for males is the July and August temperature:precipitation ratio. Floral abundance is specific to each bumble bee species and life stage and includes all flowers that could affect the production of a life stage (see Materials and Methods for details). Abundance of the previous life stage for overwintered queens is the abundance of males in the previous year (t-1); for workers is overwintered queens in the current year (t); and for males is workers in the current year (t).

**Figure 4.**
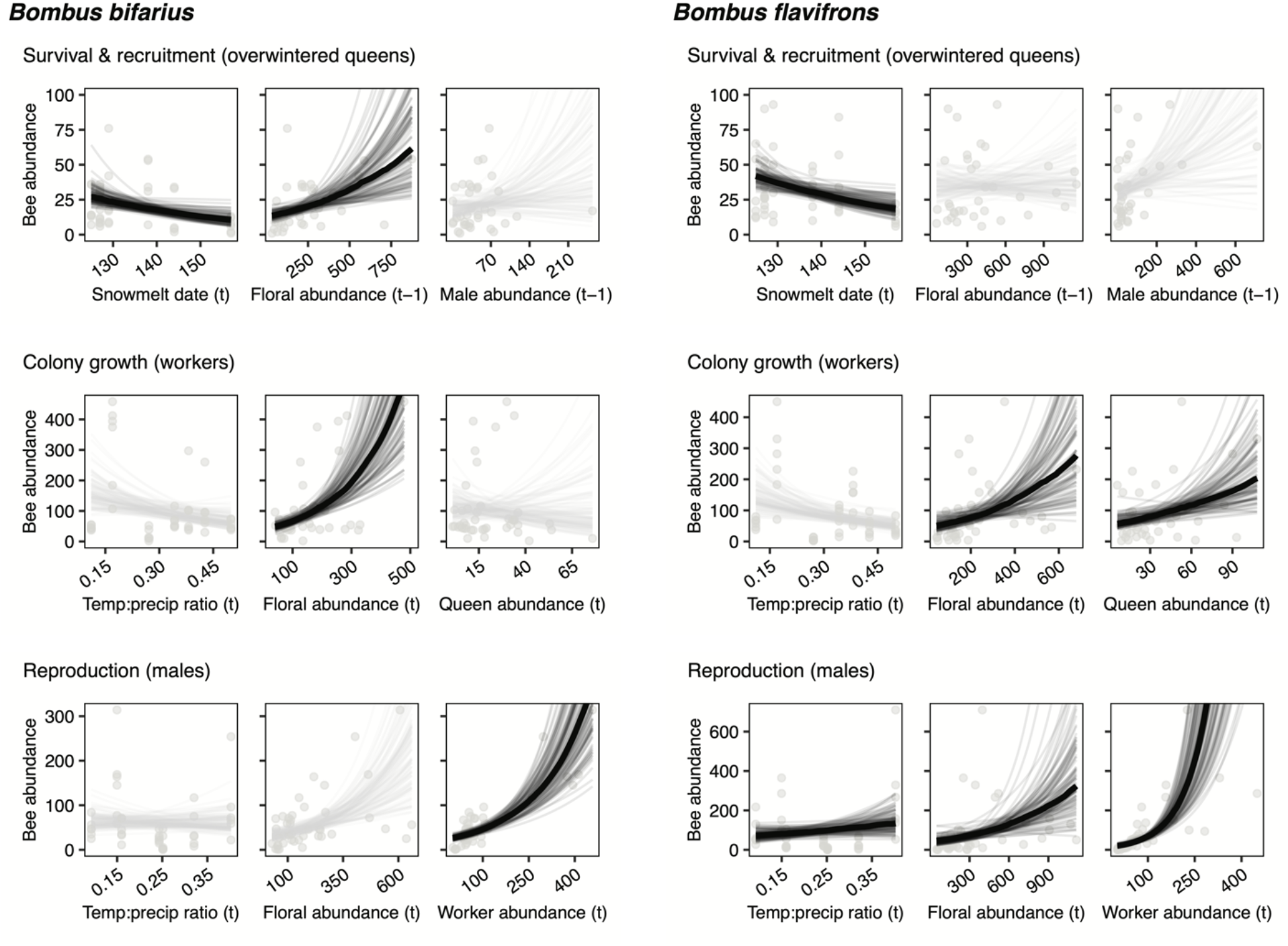
Examples of the effects of climate conditions, floral resources, and previous life stage on the abundance of different life stages for *Bombus bifarius* and *B. flavifrons*. Model fit lines represent 100 draws from the posterior distribution from a bivariate negative binomial model of each predictor variable on each response variable. Darker lines indicate evidence of a non-zero effect from the full additive model (see Fig. 3 and Materials and Methods for details) and the thick solid line represents the median estimate from the posterior distribution. Each dot represents a year-site abundance value; *n* = 36 for overwintered queens; *n* = 42 for workers and males. Similar bivariate plots for all other species are included in Supporting Information Figs. S1–5. Note that the coefficients from the full models (Fig. 3) represent effects when the other predictor variables are held constant and therefore may differ somewhat from the bivariate relationships shown here for illustrative purposes.

For the abundance of overwintered queens, our models revealed consistent evidence of a negative response to winter climate conditions (quantified as the timing of spring snowmelt) across species (5 out of 7 species; Fig. 3; Table S1). That is, more queens were observed in years with earlier snowmelt dates. For *B. bifarius*, the abundance of overwintered queens also responded to abundance of flowers in the year before queens emerged from winter diapause (t-1), showing an idiosyncratic response (Figs. 3, 4). Two species (*B. insularis* and *B. appositus)* also exhibit an effect of abundance of the previous life stage (reproductive output in year t-1 predicting overwintered queen abundance in year t; worker abundance in year t-1 showed broadly similar results, Fig. S6), although these patterns were in opposite directions, positive and negative, respectively.

The abundance of workers responded most consistently to floral resource availability, with a positive effect in four of the five remaining species, including in *B. bifarius* and *B. flavifrons* (Fig. 4); only in *B. occidentalis* was this effect not evident (Fig. 3; recall that *B. insularis*, a bumble bee nest parasite, lacks a worker life stage). We also observed a positive effect of the abundance of the previous life stage (overwintered queens) on worker abundance for *B. flavifrons*, *B. occidentalis*, and *B. rufocinctus*. Summer climate conditions (temperature:precipitation ratio for June and July) did not obviously relate to worker abundance for any species.

The abundance of males responded most consistently to the abundance of the previous life stage: worker and male numbers were positively related for all five remaining species (again *B. insularis* is not included), including in *B. bifarius* and *B. flavifrons* (Figs. 3, 4). Additionally, in all but one of these species, other factors also played a role in affecting male abundance. We observed positive effects of floral resource abundance on male abundance (4 of 7 species), and positive effects of climate conditions (3 out of 7 species), that is, we observed more males when it was warmer and drier; both factors were positively related to male abundance in *B. flavifrons* (Fig. 4).

## 4 DISCUSSION

Organisms living in seasonal environments are exposed to different sets of abiotic and biotic conditions as they transition from one life stage to the next across their life cycle. How different stages of an organism’s life cycle respond to these varying conditions is a basic feature of their biology and is critical for understanding how they will respond to environmental change. In our long-term examination of annual bumble bees, we observed a consistent set of responses in the abundance of each life stage to climate conditions, floral resource availability, and previous life stage abundance. Across the seven bumble bee species we studied, these three factors each influenced a distinct life stage, illustrating how the relative importance of different environmental factors can shift throughout pollinator life cycles (Fig. 3). The consistency of responses across species and their life stages emerges despite interspecific variation in flight season, body size, and other aspects of morphology, behaviour, and natural history, suggesting common underlying mechanisms.

The shifting importance of different abiotic and biotic factors through the bumble bee life cycle provides insight into the demographic mechanisms that may underlie how these pollinator populations respond to environmental change. Recognizing this complexity is especially relevant in an era of climate change and pollinator declines. For example, changing climate (e.g., increasing temperatures) and reductions in floral resources are both hypothesised to have strong and negative effects on many bumble bee species (Woodard & Jha, 2017; Soroye et al., 2020; Cameron & Sadd, 2020). Our results illustrate that changes in climate do not equally influence each life stage—certain stages can be more sensitive to climate conditions than others. Similarly, although floral resources are at the core of the plant-pollinator mutualism, we find their strongest effects on only some parts of the life cycle. Here, the positive effects of floral resources on worker numbers also translate into positive effects on the output of reproductive bees because the abundances of these two life stages vary consistently and positively. Other life stages were not consistently linked, however, which means that as different factors affect each life stage, there is the potential for demographic vital rates to become decoupled in their responses to environmental change.

The abundance of overwintered queens, who establish colonies after surviving diapause, responded most consistently to variation in winter climate conditions—more queens with earlier snowmelt—and generally not to floral resources or the abundance of the previous life stage. In our subalpine ecosystem queens spend 8–9 months underground in winter diapause, which is at the longer end of the known range for bumble bees (6–9 months; Alford, 1969). Earlier springs occur after shorter winters of lower snowfall, meaning that diapausing queens spend less time dependent on their lipid and carbohydrate stores, which should improve overwinter survival (Beekman et al., 1998). This survival effect may override any positive effect of floral resources or reproductive output in the previous year. On the other hand, survival of queens in laboratory settings was strongly related to nutrient acquisition before diapause (Woodard et al., 2019; Treanore & Amsalem, 2020), and overwintered queen abundance was related to floral resources in the preceding season in some field studies (Inari et al., 2012; Carvell et al., 2017). Consistent with this, overwinter success was positively related to previous-year resource availability—in addition to snowmelt timing—for two of our study species, *B. appositus* and *B. bifarius*. In our system of relatively long and harsh winters, however, winter conditions are likely to be the dominant driver of queen numbers overall.

The abundance of workers was positively associated with floral resource abundance in all of our study species except *B. occidentalis*, as also reported for other bumble bees (e.g., Pelletier & McNeil, 2003; Crone & Williams, 2016; Mola et al., 2020). More resources allow colonies to produce more workers, thereby increasing their potential to produce more reproductives in turn. Furthermore, the absence of any relationship between summer climate conditions (temperature and precipitation) and worker abundance suggests that the variation experienced over our study is within tolerable limits for colony growth. Most of our study species nest below ground (Williams et al., 2014), which should buffer colonies from extreme summer temperatures. That the previous life stage predicts worker abundance in only half our species suggests that the survival and establishment of queens need not be related to worker production. Successful colony establishment by overwintered bumble bee queens appears to be low in the wild (Goulson, 2010), which often may decouple that stage from colony growth later in the favourable growing season. However, for three species (*B. flavifrons, B. occidentalis*, and *B. rufocinctus*) we did observe positive covariation of workers with the abundance of spring queens. This may suggest that queens of these species have relatively high colony establishment rates, such that favourable conditions for them translate into positive effects on colony growth.

Finally, the abundance of male bees exhibited consistent and positive covariation with the abundance of workers in all of our species. This agrees with the intuitive premise expressed above that resource-dependent colony growth and size predict eventual reproductive output (e.g., Carvell et al., 2015; Herrmann et al., 2017; Hemberger et al., 2020). Indeed, we saw that reproductive output was positively affected by floral resource abundance for *B. appositus, B. flavifrons*, and *B. rufocinctus*. Reproductive output also increased with warmer and drier summer conditions for *B. flavifrons, B. insularis*, and *B. mixtus*, consistent with a pattern reported for another bumble bee species (Zaragoza-Trello et al., 2021; but see Holland & Bourke, 2015). Although extreme temperatures associated with climate change may adversely affect bumble bees (Iserbyt & Rasmont, 2012; Martinet et al., 2021), temperatures experienced in our system were likely within a favourable range for foraging activity (Kenna et al., 2021). We note that our estimate of reproductive output does not capture autumn queens—a life stage for which we have very limited data. However, the increased abundance of male bees with higher worker abundance suggests an increased investment in reproduction which should also increase the production of autumn queens (e.g., Herrmann et al., 2017).

Although our 7-year study encompassed considerable variation in abiotic and biotic conditions, more years of data are necessary to clarify some relationships, and indeed, some patterns may shift qualitatively with a longer temporal perspective (e.g., Thomson, 2019). For example, we expect the negative relationship between the abundance of *B. appositus* overwintered queens and reproductive output in the previous year to become positive or disappear with more years of study, because the inclusion of 2021 data reversed a previously weakly positive relationship (2021 had the highest queen numbers for *B. appositus* while reproductive output was especially low in 2020; Fig. S1). We do expect many relationships reported here to remain qualitatively (if not quantitatively) similar, so long as climate conditions remain within the range of variation we witnessed (which approximates that of the last 50 years, e.g., Cordes et al. 2021). But this is far from certain: coming decades in our study region are predicted to become increasingly warmer and drier with a greater frequency of extreme events (IPCC, 2021). As such, nonlinear or threshold responses (sensu Iler et al., 2013) might occur as organisms encounter conditions outside the historical range (e.g., CaraDonna et al., 2018; Martinet et al., 2021).

Predicting population outcomes under environmental change requires not only an understanding of how different factors influence each part of an organism’s life cycle, but also which life stages—and thus vital rates—most strongly influence population growth (e.g., Crouse et al., 1987; Caswell, 2001). Climate change in our study region is resulting in less winter snowpack and earlier spring snowmelt timing, which leads to shorter winters and fewer floral resources produced during the growing season (Ogilvie et al., 2017; Cordes et al., 2020). Whereas shorter winters appear to favour the survival of overwintered queens, as we found, these same conditions are unfavourable for colony growth and reproduction because lower input of snowmelt water and longer growing seasons tend to reduce flower densities (see Ogilvie et al., 2017). Despite these contrasting patterns, however, the lack of covariation between the abundance of workers and overwintered queens for half of our species, and the lack of covariation between overwintered queen numbers and reproductive males in the previous year, suggests that the positive and negative effects experienced by each life stage often remain isolated to that stage. Although successful establishment of colonies by overwintered queens in spring is likely to be the critical event for bumble bee population dynamics (Woodard et al., 2019), it remains unclear whether the benefits of warmer and shorter winters for queen survival will compensate for the expected negative effects of reduced floral resources on colony growth and reproduction.

Many bumble bee species are declining worldwide (Cameron & Sadd, 2020), and our study provides insight on drivers that may contribute to population declines. Among our focal species, *B. occidentalis* is of greatest conservation concern as it was once a common and widespread species in Western North America that has declined rapidly across its range over the last two decades (Cameron et al., 2011; Graves et al., 2020). At broad geographic scales, declines in *B. occidentalis* populations, like in other bumble bees, are potentially driven by a combination of stressors: climate change, parasites, pesticides, and habitat loss. In our subalpine study system, which is mostly unaffected by these stressors except for climate change, *B. occidentalis* abundance was affected by climate conditions, floral resources, and previous life stage across its life cycle. Over the 7-year study, overwintered queen abundance increased with earlier snowmelt dates, workers increased with the abundance of overwintered queens, and males increased with both worker and floral abundance (Fig. 3; Fig. S4). Given that two of the life stages are linked, effects during specific life stages can precipitate to others, highlighting that conservation efforts should consider how multiple factors play out across the life cycle.

Our long-term study demonstrates that different demographic parameters throughout the bumble bee life cycle show distinct responses to variation in the abiotic and biotic environment. Furthermore, the life-cycle approach we have taken with an assemblage of coexisting bumble bees illustrates that important details about pollinator demography are overlooked without consideration of life stage-specific responses. Ultimately, it is life-stage specific responses that shape population outcomes, not only for bumble bees and other animal pollinators, but for many other organisms living in seasonal environments.

## ACKNOWLEDGEMENTS

We thank the many people who have assisted with field work over the years: Alyssum Cohen, Vicky DeLira, Kaitlin Griffith, Emilie Gitter, Anna Hennage, Jeremy Jonas, Mikaela Jones, Kyle Spells, Valerie Watson, Daniele Wiley, and especially Justin Bain, Jackie Fitzgerald, and Gwen Kirschke. We thank William Petry and Amy Iler for advice on statistical analyses, and Nick Waser, Amy Iler, John Mola, and the Iler and CaraDonna Lab Group for constructive feedback on previous manuscript drafts. We also thank billy barr for collection of long-term climate data, Jennie Reithel for logistical support at the RMBL, and the Crested Butte Mountain Resort for access to a field site. Funding was provided in part by the RMBL, the Chicago Botanic Garden, Northwestern University, and the Western North American Naturalist. We acknowledge that our field research was conducted on the ancestral and unceded land of the Tabeguache band of the Núu-agha-tuvu-pu (Ute) people.

## DATA AVAILABILITY STATEMENT

Data will be available once accepted for publication on the Open Science Framework.

## AUTHOR CONTRIBUTIONS

J.E.O., P.J.C. and many field assistants collected the data; J.E.O. established the long-term monitoring project; J.E.O. and P.J.C. conceived the study, analysed the data, and wrote the manuscript.

## CONFLICT OF INTEREST

The authors declare no conflict of interest.

## Notes

### Competing Interest Statement

The authors have declared no competing interest.

