## Supporting Information for "The shifting importance of abiotic and biotic factors across the life cycles of wild pollinators"

**Table S1.** Full model results from negative binomial mixed effects models examining the effects of climate, floral abundance, and previous life stage abundance on the abundance of each bumble bee life stage for seven species over 7 years. Model estimates (i.e., the effect of each factor) from the negative binomial models are expressed as the change in the log count of bees per 1 unit change in the predictor variable, except for flowers, which are shown as the change in the log count of bees per 100 flowers. $R^2$ values are Bayesian $R^2$ values calculated for each full model using residual variance following Gelman et al., (2019); values shown here are the median from the posterior distribution (see Fig. 3 for 89% credible intervals). Note that for each species and life stage the duplicated $R^2$ values represent the model fit for the entire model (i.e., all three model terms).

| | Model term | R^2 | Estimate (coefficient) | SE | 89% credible interval | n |
|---|---|---|---|---|---|---|
| Queens | | | | | | |
| B. appositus | climate | 0.370 | -0.0423 | 0.0132 | [-0.0636, -0.0214] | 36 |
| B. appositus | previous.life.stage | 0.370 | -0.0219 | 0.0103 | [-0.0392, -0.0058] | 36 |
| B. appositus | flowers | 0.370 | 0.1810 | 0.1147 | [ 0.0045, 0.3615] | 36 |
| B. bifarius | climate | 0.603 | -0.0315 | 0.0106 | [-0.0481, -0.0141] | 36 |
| B. bifarius | previous.life.stage | 0.603 | 0.0025 | 0.0036 | [-0.0029, 0.0083] | 36 |
| B. bifarius | flowers | 0.603 | 0.1144 | 0.0644 | [ 0.0198, 0.2279] | 36 |
| B. flavifrons | climate | 0.568 | -0.0293 | 0.0096 | [-0.0452, -0.0140] | 36 |
| B. flavifrons | previous.life.stage | 0.568 | 0.0004 | 0.0009 | [-0.0011, 0.0018] | 36 |
| B. flavifrons | flowers | 0.568 | -0.0202 | 0.0350 | [-0.0805, 0.0337] | 36 |
| B. insularis | climate | 0.481 | -0.0464 | 0.0175 | [-0.0750, -0.0178] | 36 |
| B. insularis | previous.life.stage | 0.481 | 0.0056 | 0.0027 | [ 0.0012, 0.0097] | 36 |
| B. insularis | flowers | 0.481 | 0.1541 | 0.1105 | [-0.0337, 0.3357] | 36 |
| B. mixtus | climate | 0.499 | -0.0129 | 0.0241 | [-0.0487, 0.0263] | 36 |
| B. mixtus | previous.life.stage | 0.499 | 0.0808 | 0.0507 | [ 0.0010, 0.1719] | 36 |
| B. mixtus | flowers | 0.499 | 0.0325 | 0.1224 | [-0.1727, 0.2290] | 36 |
| B. occidentalis | climate | 0.345 | -0.0682 | 0.0285 | [-0.1142, -0.0225] | 36 |
| B. occidentalis | previous.life.stage | 0.345 | -0.0021 | 0.0567 | [-0.0908, 0.0928] | 36 |
| B. occidentalis | flowers | 0.345 | -0.0485 | 0.1357 | [-0.2716, 0.1739] | 36 |
| B. rufocinctus | climate | 0.446 | -0.0022 | 0.0178 | [-0.0301, 0.0263] | 36 |
| B. rufocinctus | previous.life.stage | 0.446 | -0.0011 | 0.0129 | [-0.0219, 0.0204] | 36 |
| B. rufocinctus | flowers | 0.446 | 0.0492 | 0.0881 | [-0.1025, 0.1936] | 36 |

***Table S1*** *continued.*

| | Model term | R^2 | Estimate (coefficient) | SE | 89% credible interval | n |
|---|---|---|---|---|---|---|
| Workers | | | | | | |
| B. appositus | climate | 0.561 | 0.5735 | 1.5448 | [-1.9647, 3.0022] | 42 |
| B. appositus | previous.life.stage | 0.561 | 0.0230 | 0.0168 | [-0.0041, 0.0500] | 42 |
| B. appositus | flowers | 0.561 | 0.3341 | 0.1295 | [ 0.1130, 0.5485] | 42 |
| B. bifarius | climate | 0.474 | 0.9589 | 1.3079 | [-1.1915, 3.1181] | 42 |
| B. bifarius | previous.life.stage | 0.474 | -0.0153 | 0.0102 | [-0.0318, 0.0023] | 42 |
| B. bifarius | flowers | 0.474 | 0.6542 | 0.1983 | [ 0.3607, 1.0027] | 42 |
| B. flavifrons | climate | 0.435 | 0.1589 | 1.5367 | [-2.2319, 2.6877] | 42 |
| B. flavifrons | previous.life.stage | 0.435 | 0.0138 | 0.0065 | [ 0.0034, 0.0250] | 42 |
| B. flavifrons | flowers | 0.435 | 0.3488 | 0.1275 | [ 0.1591, 0.5729] | 42 |
| B. mixtus | climate | 0.504 | -2.8637 | 2.1445 | [-5.998, 0.7970] | 42 |
| B. mixtus | previous.life.stage | 0.504 | 0.0109 | 0.1106 | [-0.165, 0.1906] | 42 |
| B. mixtus | flowers | 0.504 | 0.9735 | 0.4349 | [ 0.317, 1.6844] | 42 |
| B. occidentalis | climate | 0.457 | -0.4986 | 2.9091 | [-5.1930, 4.2269] | 42 |
| B. occidentalis | previous.life.stage | 0.457 | 0.4294 | 0.1839 | [ 0.1477, 0.7395] | 42 |
| B. occidentalis | flowers | 0.457 | 0.1463 | 0.2551 | [-0.2735, 0.5588] | 42 |
| B. rufocinctus | climate | 0.597 | -3.1717 | 1.9923 | [-6.4023, 0.0860] | 42 |
| B. rufocinctus | previous.life.stage | 0.597 | 0.0554 | 0.0311 | [ 0.0058, 0.1097] | 42 |
| B. rufocinctus | flowers | 0.597 | 0.4358 | 0.1616 | [ 0.1839, 0.7053] | 42 |

***Table S1*** *continued.*

| | Model term | R^2 | Estimate (coefficient) | SE | 89% credible interval | n |
|---|---|---|---|---|---|---|
| Males | | | | | | |
| B. appositus | climate | 0.562 | 0.9771 | 1.3403 | [-1.1379, 3.1074] | 42 |
| B. appositus | previous.life.stage | 0.562 | 0.0336 | 0.0078 | [ 0.0212, 0.0456] | 42 |
| B. appositus | flowers | 0.562 | 0.2117 | 0.1098 | [ 0.0373, 0.3914] | 42 |
| B. bifarius | climate | 0.767 | 0.6025 | 1.0817 | [-1.0332, 2.3604] | 42 |
| B. bifarius | previous.life.stage | 0.767 | 0.0051 | 0.0012 | [ 0.0032, 0.0072] | 42 |
| B. bifarius | flowers | 0.767 | 0.1041 | 0.0808 | [-0.0316, 0.2371] | 42 |
| B. flavifrons | climate | 0.562 | 2.5770 | 1.3075 | [0.4816, 4.5609] | 42 |
| B. flavifrons | previous.life.stage | 0.562 | 0.0093 | 0.0020 | [0.0061, 0.0125] | 42 |
| B. flavifrons | flowers | 0.562 | 0.1961 | 0.0545 | [0.1127, 0.2892] | 42 |
| B. insularis | climate | 0.407 | 2.3451 | 1.1841 | [ 0.5083, 4.2767] | 42 |
| B. insularis | flowers | 0.407 | 0.0606 | 0.0928 | [-0.0772, 0.2231] | 42 |
| B. mixtus | climate | 0.717 | 4.0635 | 1.9392 | [ 1.0535, 7.3390] | 42 |
| B. mixtus | previous.life.stage | 0.717 | 0.0661 | 0.0125 | [ 0.0458, 0.0881] | 42 |
| B. mixtus | flowers | 0.717 | 0.2358 | 0.1778 | [-0.0326, 0.5400] | 42 |
| B. occidentalis | climate | 0.531 | -2.3252 | 2.4766 | [-5.9648, 1.8434] | 36 |
| B. occidentalis | previous.life.stage | 0.531 | 0.0766 | 0.0316 | [ 0.0278, 0.1321] | 36 |
| B. occidentalis | flowers | 0.531 | 0.2826 | 0.1659 | [ 0.0011, 0.5695] | 36 |
| B. rufocinctus | climate | 0.509 | 1.0769 | 1.8497 | [-1.8650, 4.4238] | 42 |
| B. rufocinctus | previous.life.stage | 0.509 | 0.0127 | 0.0036 | [ 0.0073, 0.0186] | 42 |
| B. rufocinctus | flowers | 0.509 | 0.3684 | 0.1072 | [ 0.2134, 0.5566] | 42 |

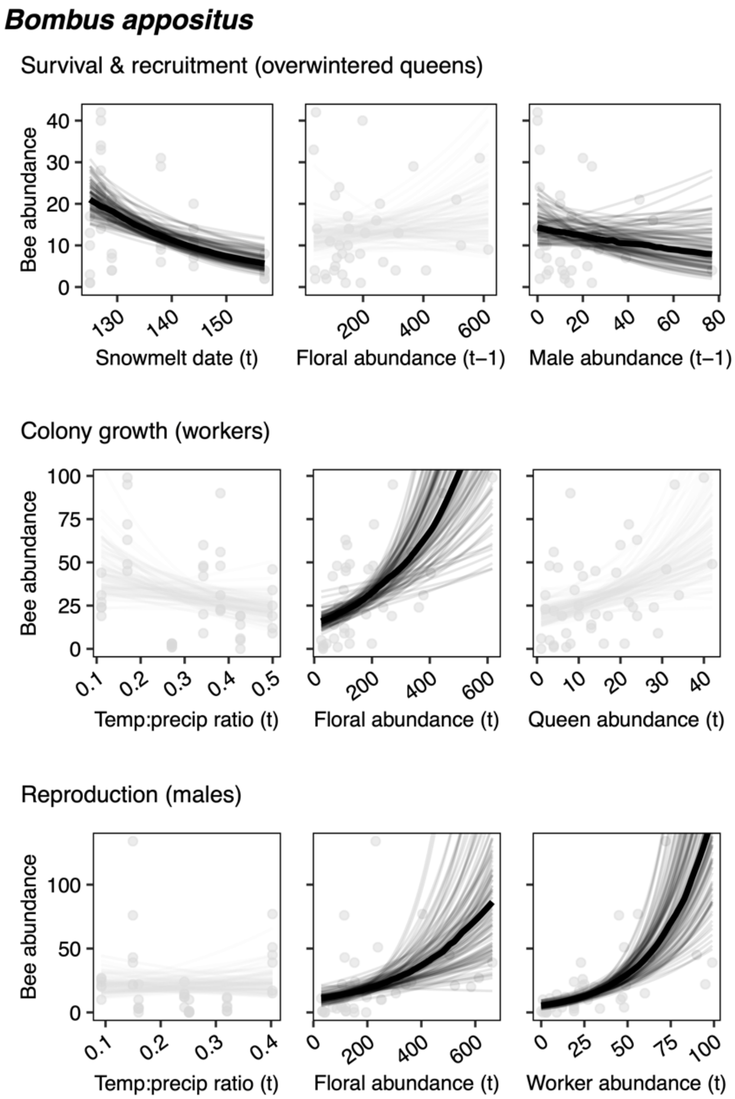


**Figure S1. The effects of climate conditions, floral resources, and previous life stage on the abundance of different life stages in *Bombus appositus*.** Model fit lines represent 100 draws from the posterior distribution from a bivariate negative binomial model of each predictor variable with each response variable. Darker lines indicate estimates from the full additive model where the 89% credible interval for the effect of each factor on bee abundance does not include zero; the thick solid line represents the median estimate from the posterior distribution (see Fig. 3 and Materials and Methods in the main text for details). Each dot represents a year-site abundance value; *n* = 36 for overwintered queens, *n* = 42 for workers and males.

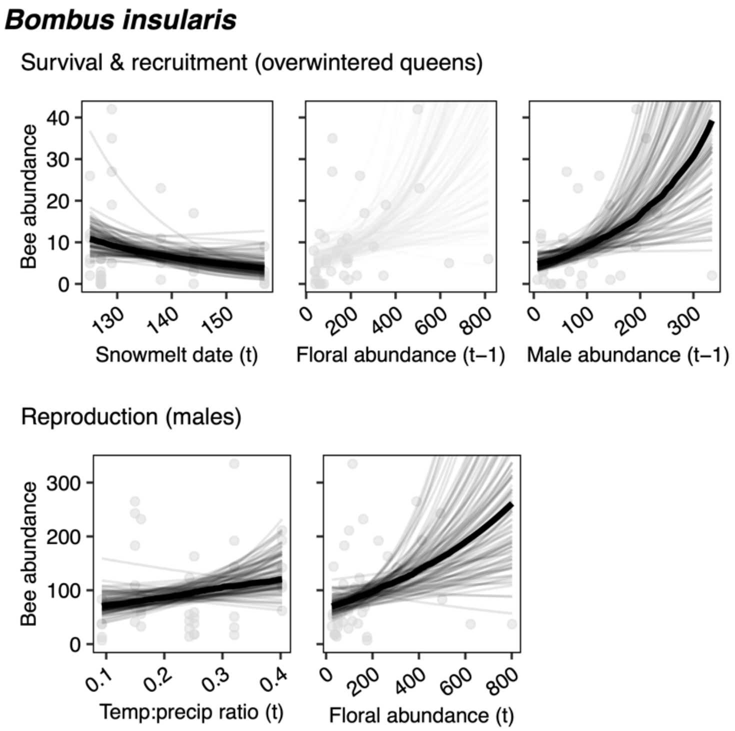


**Figure S2. The effects of climate conditions, floral resources, and previous life stage on the abundance of different life stages in *Bombus insularis*.** Model fit lines represent 100 draws from the posterior distribution from a bivariate negative binomial model of each predictor variable with each response variable. Darker lines indicate estimates from the full additive model where the 89% credible interval for the effect of each factor on bee abundance does not include zero; the thick solid line represents the median estimate from the posterior distribution. *Bombus insularis* is a bumble bee nest parasite without a worker life stage. Each dot represents a year-site abundance value; $n$ = 36 for overwintered queens, $n$ = 42 for males.

### ***Bombus mixtus***

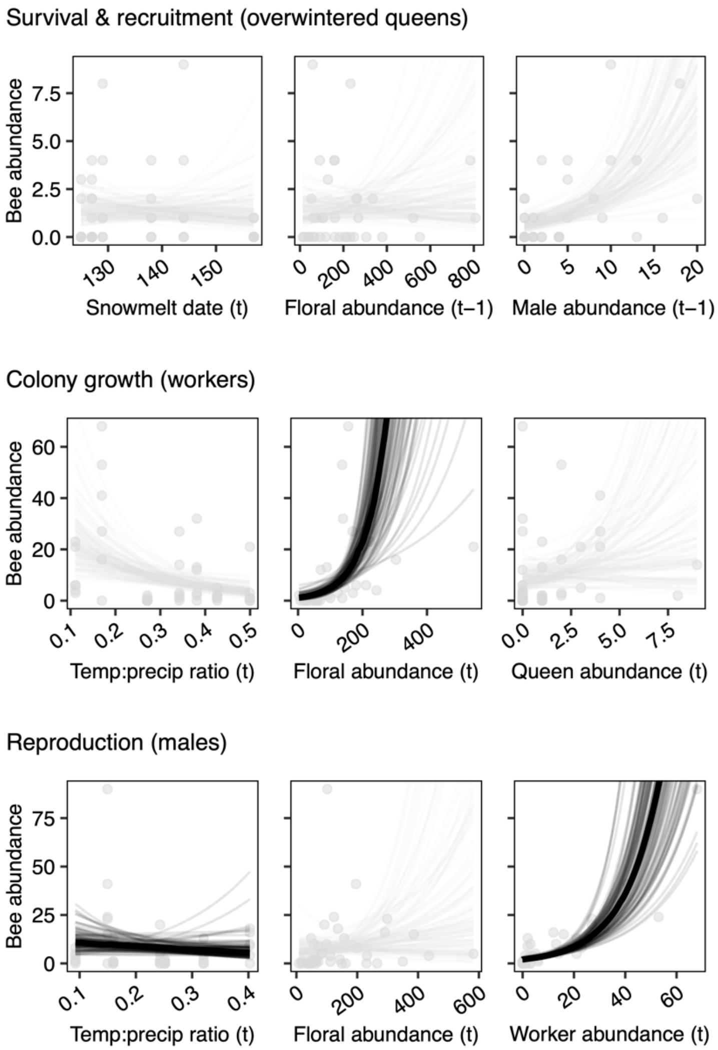


**Figure S3. The effects of climate conditions, floral resources, and previous life stage on the abundance of different life stages in *Bombus mixtus*.** Model fit lines represent 100 draws from the posterior distribution from a bivariate negative binomial model of each predictor variable with each response variable. Darker lines indicate estimates from the full additive model where the 89% credible interval for the effect of each factor on bee abundance does not include zero; the thick solid line represents the median estimate from the posterior distribution. Note that the bivariate relationship between male abundance and the temperature:precipitation ratio here appears flat, however, in the full additive model when floral abundance and worker abundance are held constant, the relationship is positive (see Fig. 3 in the main text). Each dot represents a year-site abundance value; *n* = 36 for overwintered queens, *n* = 42 for workers and males.

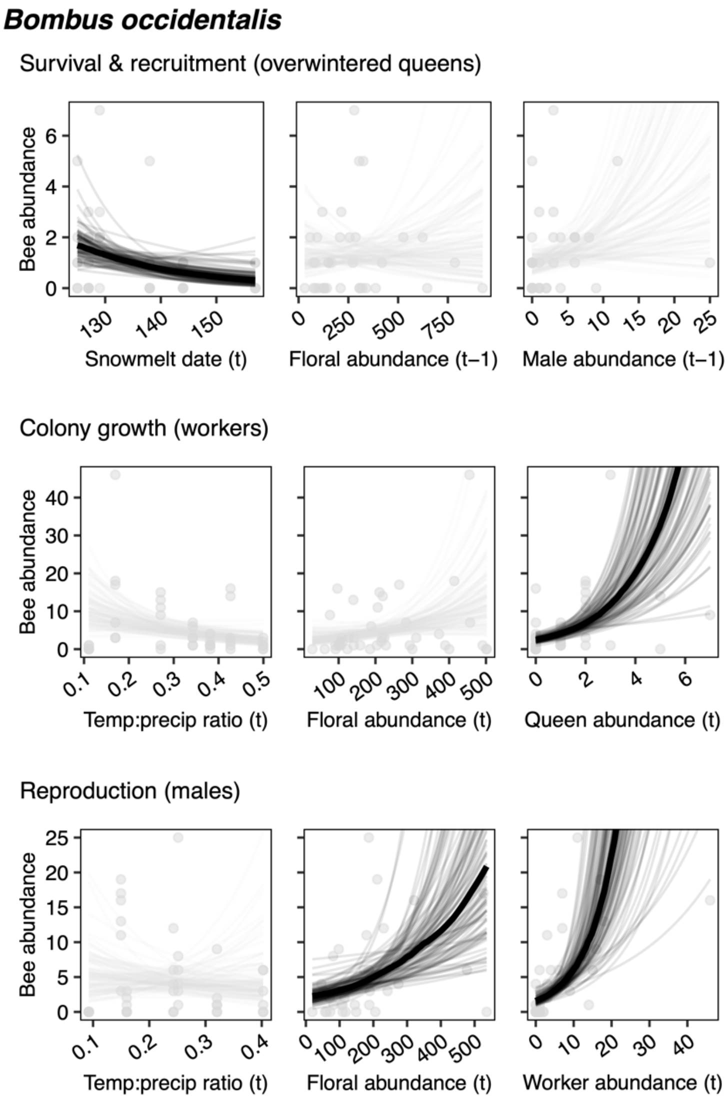


**Figure S4. The effects of climate conditions, floral resources, and previous life stage on the abundance of different life stages in *Bombus occidentalis*.** Model fit lines represent 100 draws from the posterior distribution from a bivariate negative binomial model of each predictor variable with each response variable. Darker lines indicate estimates from the full additive model where the 89% credible interval for the effect of each factor on bee abundance does not include zero; the thick solid line represents the median estimate from the posterior distribution. Each dot represents a year-site abundance value; *n* = 36 for overwintered queens and males, *n* = 42 for workers.

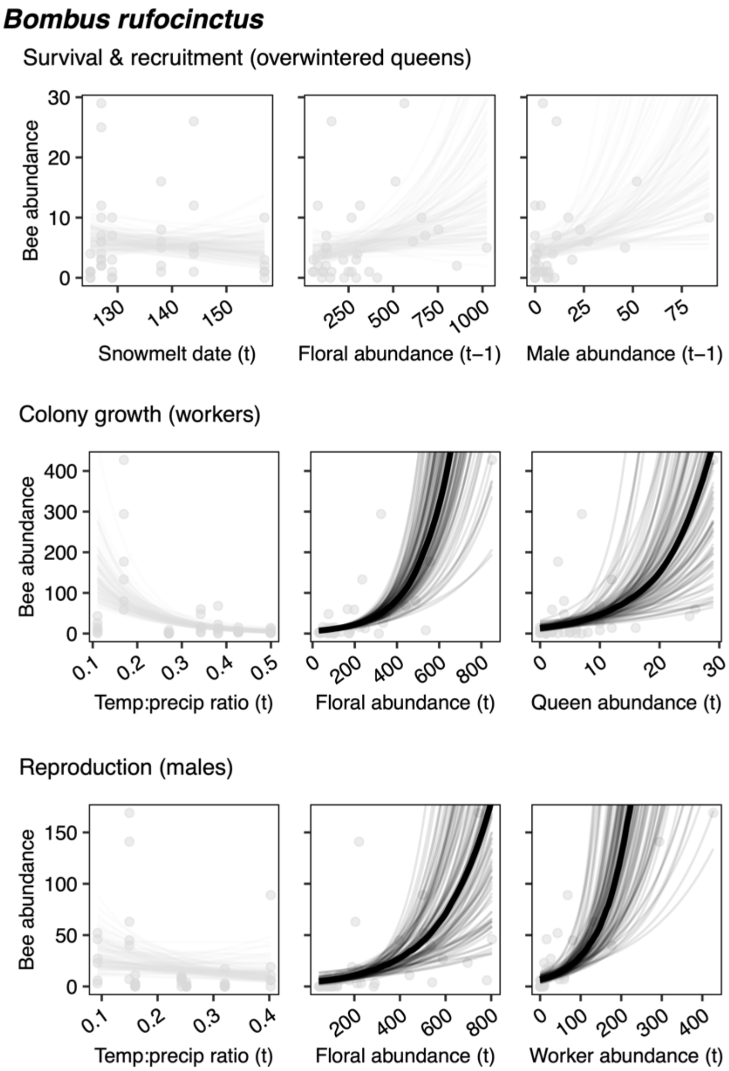


**Figure S5. The effects of climate conditions, floral resources, and previous life stage on the abundance of different life stages in *Bombus rufocinctus*.** Model fit lines represent 100 draws from the posterior distribution from a bivariate negative binomial model of each predictor variable on each response variable. Darker lines indicate estimates from the full additive model where the 89% credible interval for the effect of each factor on bee abundance does not include zero; the thick solid line represents the median estimate from the posterior distribution. Each dot represents a year-site abundance value; $n = 36$ for overwintered queens, $n = 42$ for workers and males.

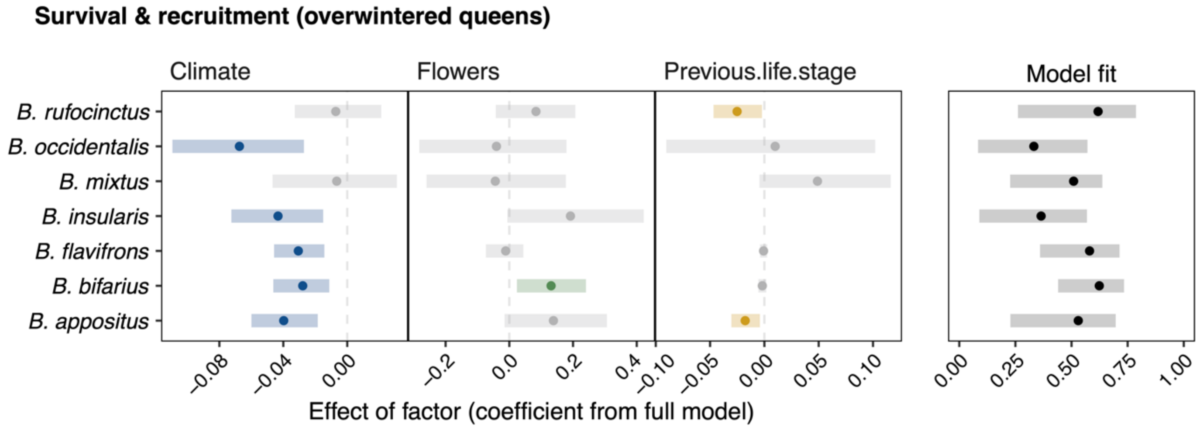


**Figure S6. The effect of snowmelt date, floral abundance, and previous life stage on the abundance of overwintered queens in the seven bumble bee species, using workers (t – 1) as the previous life stage.** Dots represent the median estimate (i.e., effect) of the posterior distribution from negative binomial mixed models including all three factors; error bars represent 89% Bayesian credible intervals. Coloured dots and error bars represent model factors with evidence in support of a non-zero effect (e.g., 89% credible interval does not include zero). Model fits represent the median and 89% credible interval for Bayesian $R^2$ values. Floral abundance is specific to each bumble bee species and includes all flowers in the previous season (see Materials and Methods in the main text for details).
